# Impact of Human Genetic Variation Underlying Resistance to Antigen-Specific Immunotherapy

**DOI:** 10.1101/2025.05.02.651174

**Authors:** Romina Marone, Erblin Asllanaj, Giuseppina Capoferri, Torsten Schwede, Lukas T. Jeker, Rosalba Lepore

**Affiliations:** Department of Biomedicine, University Hospital Basel and University of Basel, Hebelstrasse 20, CH-4031 Basel, Switzerland; Transplantation Immunology & Nephrology, University Hospital Basel, Petersgraben 4, CH-4031 Basel, Switzerland; Biozentrum, University of Basel, Spitalstrasse 41, CH-4056 Basel, Switzerland; SIB Swiss Institute of Bioinformatics, Spitalstrasse 41, CH-4056 Basel, Switzerland; Innovation Focus Cell Therapies, University Hospital Basel, Petersgraben 4, CH-4031 Basel, Switzerland

**Keywords:** antigen, antibody, immunotherapy, human genetic diversity, single nucleotide variant

## Abstract

Monoclonal antibodies have transformed the therapeutic landscape across oncology, immunology, and infectious diseases by enabling high-affinity, antigen-specific targeting, e.g. to neutralize soluble molecules, block cellular interactions or deplete cells. While high specificity is critical for safety, it may confer an inherent susceptibility to even minor variations within the target epitope. Here, we present the first systematic study investigating the impact of natural single nucleotide variants on antigen recognition by therapeutic monoclonal antibodies, both approved and in clinical development. For every antibody analysed, we identified genetic variants within or near the antibody-antigen interface, a subset of which are predicted to disrupt antigen recognition. Experimental studies corroborated the impact of select variants for four different antigens, revealing complete loss of antibody binding in some cases. As a consequence, a human breast cancer cell line overexpressing ErbB2 but engineered to carry the ErbB2^P594H^ variant, was completely resistant to killing even by highly potent clinical antibody-drug conjugates (ADCs). These findings suggest that natural variants can confer primary resistance to antibody-based therapies, with critical implications for treatment outcomes, patient management and safety, particularly in the context of potent modalities such as ADCs. Notably, individual resistance-associated variants, while globally rare, are enriched in specific populations, underscoring the importance of accounting for genetic diversity in both drug development and clinical decision-making.

**One sentence summary:** Naturally occurring genetic variants in human therapeutic target antigens can affect binding by almost any therapeutic antibody.

## Introduction

The discovery that individual lymphocytes express antigen receptors of a single specificity and that B lymphocytes secrete antibodies of a single specificity, resulted in methods to produce monoclonal antibodies (mAb) (*1–3*). The antibody repertoire in an individual human is estimated to exceed 10^12^, with theoretical diversity orders of magnitude greater, enabling the development of antibodies against a wide range of targets and making them highly attractive as diagnostic and therapeutic tools (*4*). Antibodies recognize and bind their target antigens (Ag) through the complementarity-determining region (CDR), interacting with specific regions of the antigens known as epitopes. A few amino acid residues within a given epitope are critical for the mAb-Ag interaction. Some antibodies can even discriminate differences as small as a single amino acid substitution (*5, 6*) and picomolar (pM) binding affinities can be reached. Such extraordinarily high diversity, specificity and affinity enables antigen-specific, targeted therapies against almost any antigen. Indeed, the ability to target specific antigens led to the first commercial approval of a mAb to deplete T cells in 1986 (OKT3) (*7*). Later, a CD20-directed, B cell depleting mAb became the standard of care to treat B cell lymphomas (*8*). Many other therapeutic antibodies followed and antibody engineering and development enabled a major industry (*9*). Today, >100 antibody-based therapeutics are approved for many different diseases (*9*). Thus, the development of diagnostic and therapeutic mAbs revolutionized biology and medicine (*2, 9, 10*).

Therapeutic antibodies can exert their function through immune-mediated mechanisms such as antibody-dependent cellular cytotoxicity (ADCC), antibody-dependent cellular phagocytosis (ADCP), complement-dependent cytotoxicity (CDC), or can be used as a vehicle to deliver e.g. a toxic payload to a specific target (antibody-drug conjugate, ADC). If a purely blocking antibody is desired, unwanted natural immunologic functions can be removed by antibody engineering (e.g. silencing the Fc region), demonstrating the versatility of antibodies. Examples of therapeutic antibodies using these modes of action (MoA) are rituximab (CD20, ADCC), trastuzumab emtansine and trastuzumab deruxtecan (CD340 (ErbB2 aka HER2), ADC) and nivolumab and pembrolizumab (CD279 (PD-1), blockade), respectively (*8, 11–13*). Furthermore, antibodies can be engineered to have dual (or higher) specificity which can be used to engage T cells to kill a specific target cell (*10, 14, 15*). In addition, the antigen binding-domain can be engineered to increase binding affinity to the antigen and/or it can be fused to cellular receptors as single-chain variable fragments (ScFvs), resulting in chimeric antigen receptor (CAR) cells (*16*). Today, >7 CAR products are commercially available and >30’000 patients have been treated, a number that is rapidly increasing (*17*). Many other advanced antibody-based therapeutic formats are being developed, as recently reviewed (*9, 10*). Collectively, millions of patients are treated with antibody-derived products every year, improving health, saving many lives and representing an important economic sector (*10*).

Based on the observation that mAbs can discriminate epitopes that differ by a single amino acid (*6*), we have demonstrated that substituting those key residues using genome engineering abolishes mAb binding to the successfully engineered cells (*18*). Specifically, we and others demonstrated that designed single amino acid substitutions are sufficient to completely abolish binding and consequently cytotoxicity mediated by ADCC, T cell engagers (TCE) and even highly potent ADCs or CAR T cells for human CD123, CD117 and CD45, respectively (*19–23*). Therefore, rendering therapeutic cells resistant to antibody-derived therapy can be used to create a synthetic selectivity for diseased cells and to widen the therapeutic window. Collectively, epitope engineering is emerging as a new promising strategy with potential applications to treat leukemias, reduce toxicity in hematopoietic stem cell transplantation, and to treat HIV (*19–23*). Remarkably, we noted that single amino acid substitutions that achieve complete shielding include naturally occurring variants. For instance, E51K results from a naturally occurring single nucleotide variant (SNV) in CD123, the interleukin-3 alpha chain, that completely abolishes binding of the high affinity mAb CSL362 (*20*). Based on these findings, and considering the rapid expansion of antibody-based therapies in terms of both therapeutic formats and patient populations (*10*), we hypothesized that collectively, human genetic variation could impact the clinical efficacy of antigen-specific immunotherapies, even though individual variants may be rare. Indeed, a literature search confirmed individual case studies. For instance, a single missense mutation in the complement protein C5 results in a poor response to treatment with eculizumab (*24*). Here, we systematically analyze the prevalence of naturally occurring SNVs across human therapeutic antigens and examine how the resulting protein variants may affect the binding of therapeutic antibodies, both approved and in clinical development. Remarkably, for every therapeutic antibody analyzed, we identified at least one SNV that could impair antibody binding, and consequently, affect clinical efficacy. Our results could have important implications for patients, physicians and the pharmaceutical industry.

## Results

### Prevalence of natural missense variants across monoclonal antibody epitopes

The dataset analyzed in this study includes 87 distinct antibody sequences (mAbs), encompassing a wide range of clinical stages, molecular targets, and therapeutic areas (Fig. 1A and Data file S1). Among these, 44 mAbs are approved for clinical use, while the remaining are in various phases of clinical development (Phase I–III). The therapeutic areas represented include oncology (40 mAbs), immunology (28 mAbs), hematology (8 mAbs), neurology (7 mAbs), and other indications (4 mAbs), such as infectious disease, pain management, musculoskeletal and eye disorder. The antibodies target 62 distinct antigens, including key targets such as the Programmed cell death protein 1 (PD-1) and its ligand (PD-L1), the B lymphocyte antigen CD20, the receptor tyrosine-protein kinase ErbB2, as well as others, with some being targeted by multiple mAbs. This diverse dataset offers a comprehensive snapshot of the current landscape of therapeutic antibody development (*25, 26*).

**Fig. 1.**
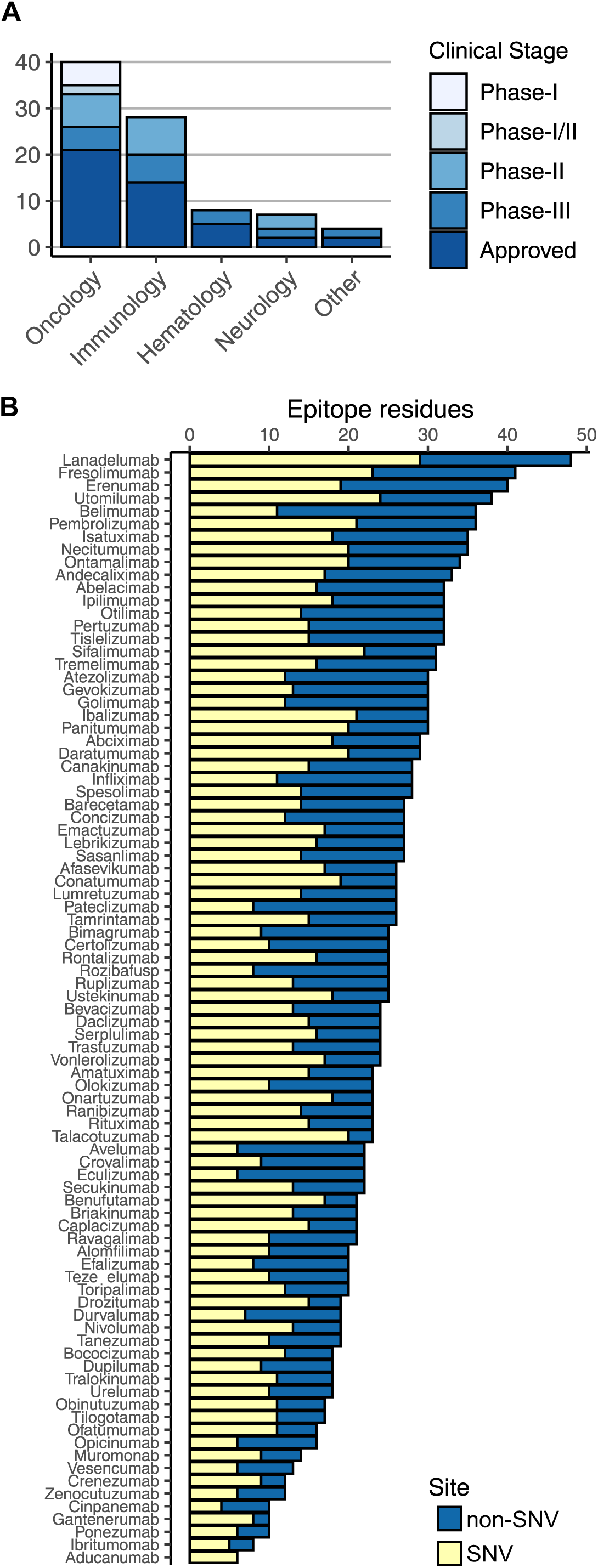
Dataset overview of therapeutic antibodies. **(A)** Distribution of antibodies by clinical stage and therapeutic indications. **(B)** Distribution of SNV sites across antibody epitopes.

For each antibody, we analyzed available 3D structures of antibody-antigen complexes to define the epitope as the set of amino acid residues at the interface of the complex. We subsequently assessed the occurrence of genetic missense variants, affecting epitope residues using a comprehensive dataset derived from population-wide studies (see Methods). Among 25,453 distinct variants associated with the therapeutic targets in our dataset, 10,352 mapped to structurally determined protein regions, of which, 1,389 corresponded to epitope regions. Notably, for every antibody in our dataset, we identified multiple missense variants within their epitope regions, with the only exception of the fresolimumab-TGFB1 epitope. All other analyzed epitopes, including those formed by fresolimumab with its alternative targets (TGFB2, TGFB3), contain multiple SNV sites (Fig. 1B). On average, approximately 50% of the residues within epitope regions are annotated with one or more missense variants based on our dataset.

### Structural and functional effects of missense variants on antigen recognition

To estimate the effect of SNV-associated amino acid variants, we employed a structure-based computational approach relying on an empirical force field to calculate the free energy change (ΔΔG) between the wildtype (WT) and the SNV mutant amino acid (*27*). Figure 2A shows the predicted ΔΔG for each variant, comparing their effect on the stability of the antibody-antigen complex (*y* axis) versus the antigen alone (*x* axis). Data points are color-coded based on the mutational impact predicted by AlphaMissense (AM) (*28*). Variants with ΔΔG > 1 (Ab-Ag complex) are predicted to impact antibody binding to the antigen, while variants with ΔΔG > 1 (Ag) are predicted to disrupt antigen stability which could indirectly affect antibody binding as well (*29*). Approximately 50% of the variants are predicted to be neutral, i.e. showing a ΔΔG ≤ 1 for both the antibody-antigen complex and the antigen (lower left quadrant). The remaining are predicted to impact stability, with ∼20% affecting antibody binding while being tolerated by the antigen (upper left quadrant), and ∼30% impacting antigen stability and antibody binding (upper right quadrant). Notably, variants predicted to impact antigen stability tend to show higher average AM scores with a substantial proportion predicted as ‘ambiguous’ or ‘likely pathogenic’ by AM (Fig. 2A and fig. S1A). To classify variants based on their predicted effect, we computed the consensus between both predictions (Fig. 2B) and plotted the resulting SNV classification for each antibody (Fig. 2C). Specifically, 41% of variants are predicted to be neutral by both methods, 10% are predicted to affect the antigen itself and 15% are predicted to interfere with antibody binding. The latter refers to variants predicted to affect the stability of the Ab-Ag complex but have a neutral effect on the antigen, according to both methods. The remaining 34% of variants are classified as uncertain, encompassing variants where the two prediction methods yield conflicting results (e.g., ΔΔG < 1 and likely pathogenic by AM), as well as variants for which AM predicts an ambiguous effect (fig. S1B), representing approximately one-third of the uncertain category. As expected, epitope residues associated with SNVs predicted to impact antigen structure or function are typically more buried and exhibit lower sequence variability in multiple sequence alignments (MSA) compared to other epitope residues (fig. S1, C and D). Conversely, residues associated with SNVs predicted to be neutral for the antigen or only affecting antibody binding display higher levels of solvent accessibility and sequence variability. Furthermore, SNVs predicted to impact antigen structure or function contain a higher proportion of known disease-associated variants compared to SNVs in the other three categories (fig. S1E). In summary, 84 out of the 87 antibodies in our dataset bind to epitopes for which we identify multiple SNVs predicted to either impact antibody binding or to affect the antigen itself (Fig. 2C). Among these, we accurately predict the effects of variants known to abolish antibody binding, such as C5^R885H^, which causes therapy failure with eculizumab (*24*), and CD38^S274F^, which disrupts daratumumab binding (*30*). Furthermore, we correctly predict the effects of multiple IL3RA variants, including E51K which we previously showed to abolish talacotuzumab binding, and S59P, which impacts both talacotuzumab and IL-3 binding (Data File S2) (*20*).

**Fig. 2.**
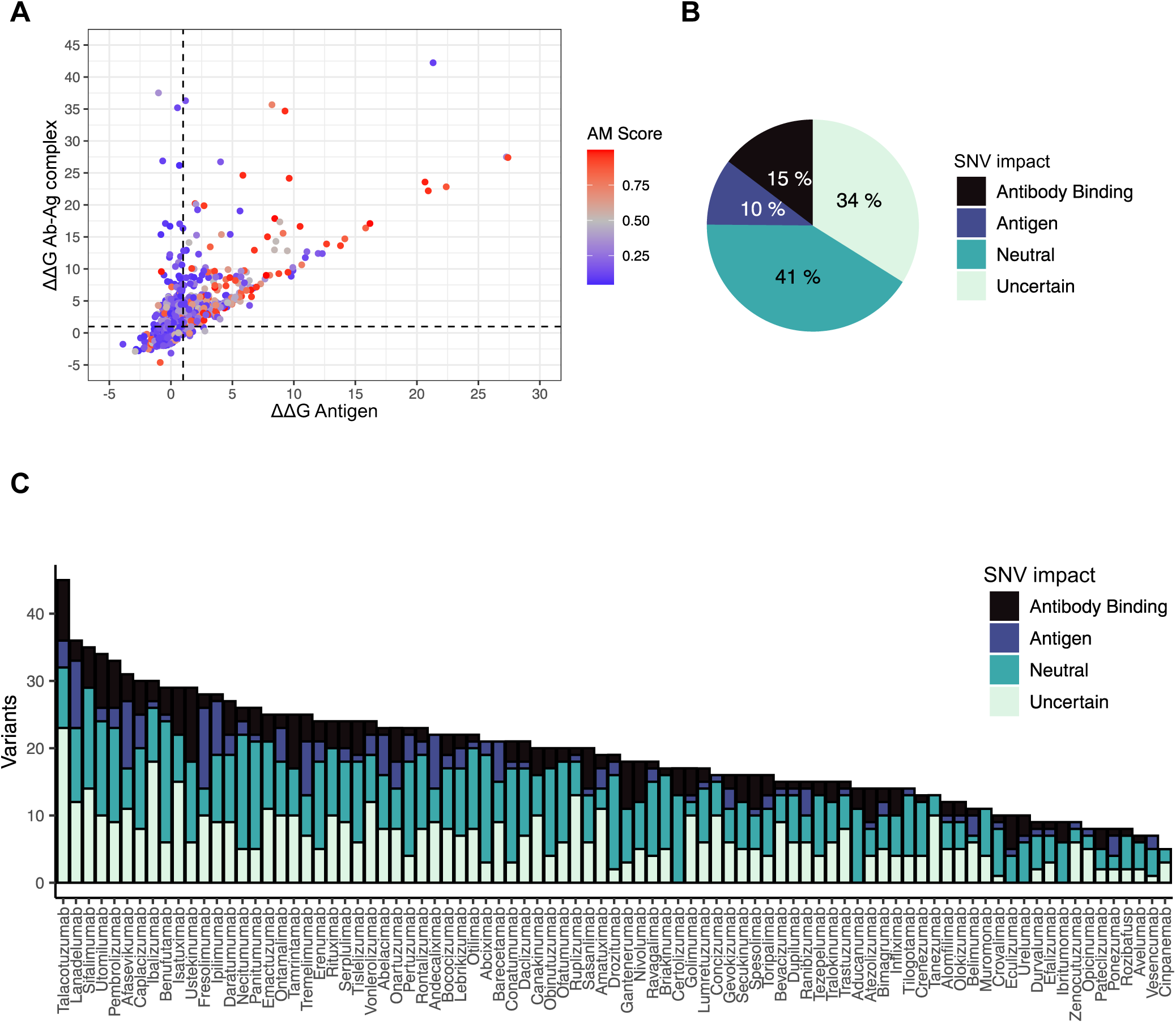
Predicted effect of missense epitope variants on antibody binding and antigen stability. **(A)** Predicted effect (ΔΔG) of missense epitope variants on antibody-antigen complexes (y-axis) versus antigens alone (x-axis). Data points are color-coded based on AlphaMissense (AM) scores. Variants with low AM scores (likely benign) are shown in blue, high scores (likely pathogenic) are shown in red, and intermediate scores (ambiguous) are shown in grey. **(B)** Classification of SNV impact based on consensus (or lack thereof) between structure-based predictions and AM scores across the full dataset and **(C)** by individual epitopes.

To investigate these results further, we selected CD20 (rituximab), CD38 (daratumumab, isatuximab), CD279 (PD-1) (nivolumab, pembrolizumab) and CD340 (ErbB2) (trastuzumab, pertuzumab) for a more in-depth characterization and experimental validation. This selection was guided by computational predictions, clinical relevance, antibody availability, and mode of action.

### Single Nucleotide Variants affecting B cell depleters

CD20 is a key drug target for B cell depletion therapy in B cell lymphomas and autoimmune diseases (*31, 32*). The introduction of CD20-targeting mAbs, such as rituximab, a very successful mAb that rapidly became the standard of care to treat B cell lymphomas (*8*), has driven the development of successive generations of anti-CD20 antibodies, e.g. ofatumumab. The epitopes of rituximab and ofatumumab are distributed across two extracellular loops (ECL1: residues 70-82 and ECL2: residues 142–182) which, according to our analysis, host multiple SNV sites (Fig. 3A). Rituximab binds to the dimeric form of CD20 in a 2:2 stoichiometry (Fig. 3B), with an interaction interface primarily mediated by ECL2 and the rituximab heavy chain CDRs (Fig. 3, B and C) (*33*). Ofatumumab, a second-generation mAb, also binds CD20 in a 2:2 stoichiometry but with a distinct geometry (Fig. 3D), with the ECL2 interacting with both the heavy and light chain CDRs (Fig. 3, D and E), resulting in more efficient complement recruitment (*34, 35*). Most identified variants are predicted to have a neutral effect on antibody binding, as well as on the structure and function of the protein (Fig. 3A), except variant N171Y. Residue N171 is in the C-terminal region of the ECL2 loop, within the ^170^ANPS^173^ motif, a critical recognition site for several anti-CD20 antibodies, including rituximab and the chimeric 2H7 mAb (*36–38*). Computational predictions suggest a differential impact of the N171Y variant on rituximab and ofatumumab binding (Fig. 3A). Analysis of the experimental structure of the Ab-Ag complex reveals that N171 directly interacts with rituximab’s heavy chain loop H3 through a hydrogen bond with S95 and van der Waals interactions with W100B (H3) and H35 (H1) (Fig. 3C). In contrast, within the CD20-ofatumumab interface, N171 primarily interacts through hydrophobic interactions with the light chain loop L3, facing residue W94. A key interaction involves the adjacent residue CD20 E174, which forms a hydrogen bond with S56 on the heavy chain loop H2 of ofatumumab (Fig. 3E). This distinct interaction pattern might explain the predicted resilience of ofatumumab to the N171Y variant.

**Fig. 3.**
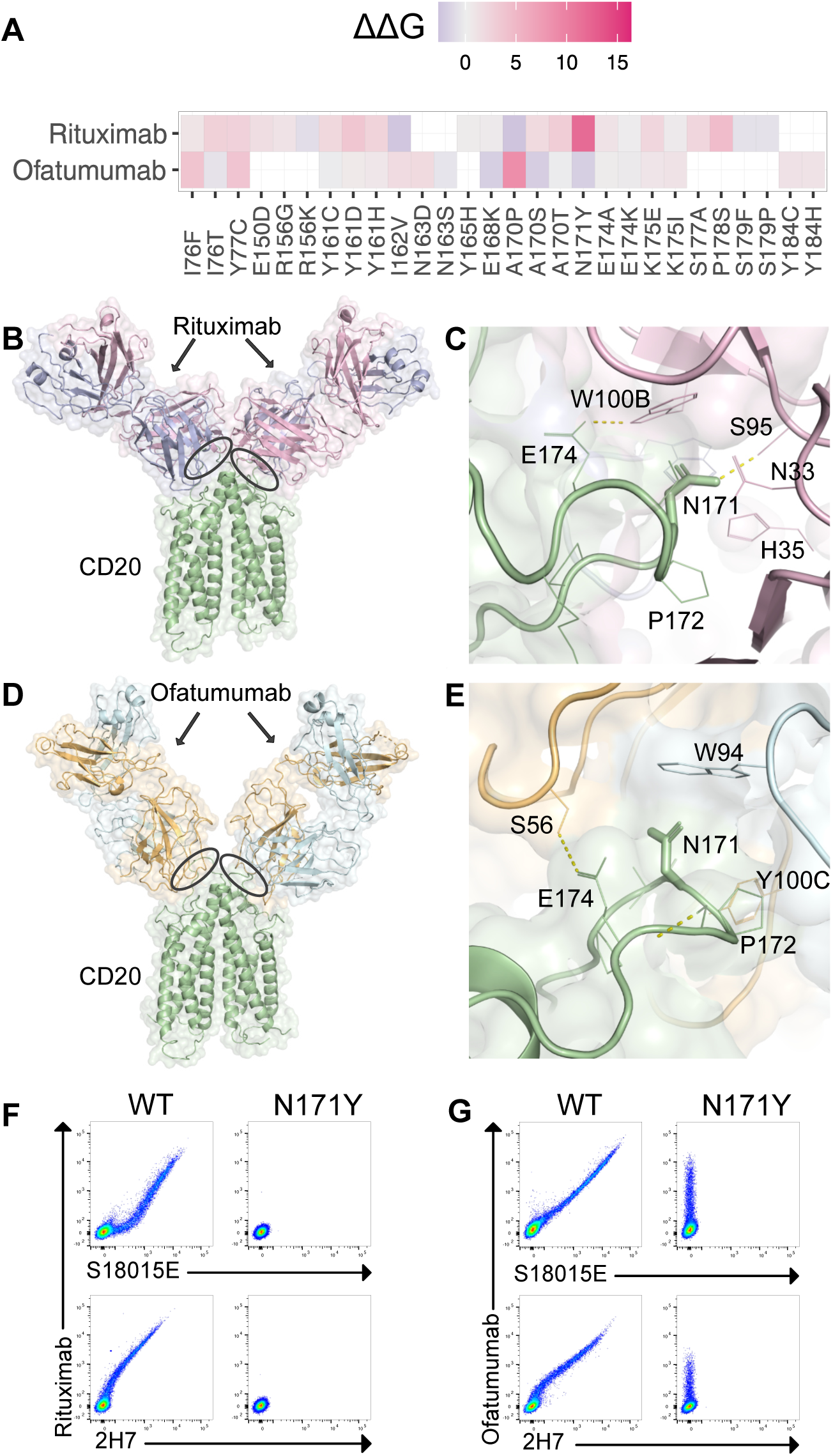
Impact of the CD20 N171Y variant on the binding of rituximab and ofatumumab antibodies. **(A)** Predicted changes in binding free energy (ΔΔG) for missense epitope variants in the CD20-rituximab and CD20-ofatumumab complexes, represented as a heatmap. **(B)** Three-dimensional structure of the 2:2 CD20-rituximab complex (PDB ID: 6Y90). The CD20 antigen is shown in green, with the rituximab heavy chain in pink and light chain in blue. Ovals indicate the locations of SNV sites at the interface of the complex. **(C)** Close-up view of the N171 residue at the CD20-rituximab interface. Residue N171 is depicted as a stick, with neighboring CD20 residues (green) and rituximab residues (pink and blue) shown as lines. Antibody residues are labeled according to the Chothia numbering scheme. Hydrogen bonds are indicated with yellow dashed lines. **(D)** Three-dimensional structure of the 2:2 CD20-ofatumumab complex (PDB ID: 6Y92). The CD20 antigen is shown in green, with the ofatumumab heavy chain in light blue and light chain in orange. Ovals indicate the locations of SNV sites at the interface of the complex. **(E)** Close-up view of the N171 residue at the CD20-ofatumumab interface. Residue N171 is depicted as a stick, with neighboring CD20 residues (green) and ofatumumab residues (light blue and orange) shown as lines. Antibody residues are labeled according to the Chothia numbering scheme. Hydrogen bonds are indicated with yellow dashed lines. **(F)** Flow cytometry histograms of CD20 WT or N171Y variant overexpressed in DF-1 cells. Various combinations of staining using mAbs rituximab, S18015E and 2H7. **(G)** Flow cytometry histograms of CD20 WT or N171Y variant overexpressed in DF-1 cells. Various staining combinations using mAbs ofatumumab, S18015E and 2H7. For (F) n = 4 independent experiments for each group were performed. For (G) n = 2 independent experiments for each group were performed.

To validate the computational predictions, we expressed CD20^WT^ and CD20^N171Y^ in chicken DF-1 cells that do not naturally express human CD20 and analyzed antibody binding by flow cytometry. Besides rituximab we used two control antibodies (clones S18015E and 2H7). We previously found that this is a sensitive assay to determine the effect of SNVs affecting mAb binding to extracellular domains (ECDs) (*19, 20*). CD20^WT^ expressing cells were double stained with rituximab and each of the control antibodies. However, CD20^N171Y^ expressing cells were not bound by rituximab and both control antibodies suggesting that CD20^N171Y^ affected binding of all three antibodies (Fig. 3F). In contrast, staining with ofatumumab was preserved, confirming the expression of the N171Y variant, while binding of the control mAbs (clones S18015E; 2H7) was abolished (Fig. 3G). Thus, in line with our computational analysis, CD20^N171Y^ specifically abolishes the binding of rituximab, antibodies S18015E and 2H7, but not ofatumumab.

### Single Nucleotide Variants impacting anti-CD38 antibodies isatuximab and daratumumab

Daratumumab and isatuximab are the two anti-CD38 mAbs currently approved for the treatment of multiple myeloma. Their interaction with CD38 triggers various downstream mechanisms, including ADCC, ADCP, and CDC (*39*), leading to effective anti-tumor activity. These mAbs target distinct CD38 epitopes (Fig. 4, A and B). Isatuximab binds to the catalytic region of CD38, exhibiting strong pro-apoptotic activity independent of cross-linking agents, unlike daratumumab (*40, 41*). Epitope regions of both antibodies are associated with multiple SNV sites (Fig. 4A). Among these, the CD38^G113R^ variant is predicted to have the most significant impact on isatuximab binding, with potential consequences on the antigen itself (Fig.4, A and C). Additionally, CD38^T148M^ and CD38^R194H^ are predicted to affect isatuximab binding, although to a lesser extent. For daratumumab, the CD38^S274F^ variant is predicted to substantially impair antibody binding, in agreement with previous findings (*30*). In contrast, variants at residue W241, which is central to a β-sheet, are predicted to impact the antigen itself (Fig.4, A and D), potentially causing local perturbations that might alter the binding of the antibody, with CD38^W241S^ showing a more pronounced effect based on predictions.

**Fig. 4.**
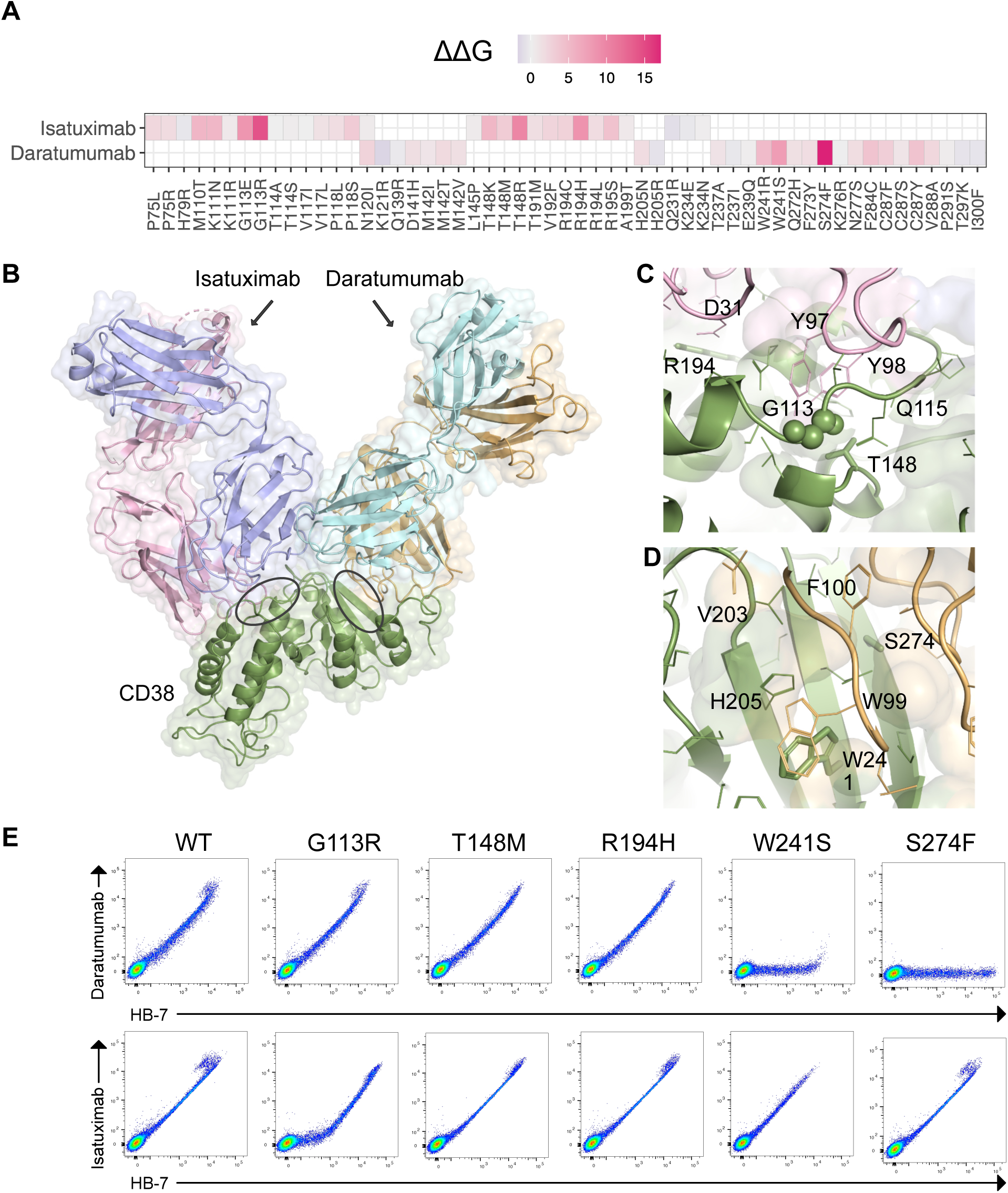
Impact of CD38 epitope variants on the binding of isatuximab and daratumumab antibodies. **(A)** Predicted changes in binding free energy (ΔΔG) for missense epitope variants in the CD38-isatuximab and CD38-daratumumab complexes, represented as a heatmap. **(B)** Three-dimensional structure representation of the CD38 ECD in complex with isatuximab (PDB ID: 4CMH) and daratumumab (PDB ID: 7DHA). The visualisation is derived from the structural superposition of CD38 (green). The isatuximab heavy chain is shown in pink, and its light chain in blue. Daratumumab heavy chain is shown light blue, and its light chain in orange. Ovals indicate the locations of SNV sites at the interface of the complex. **(C)** Close-up view of key epitope residues (G113, T148 and R194) at the CD38-isatuximab interface. The residues are shown as sticks (G113 main chain atoms as spheres), with neighboring CD38 residues (green) and isatuximab residues (pink and blue) shown as lines. Antibody residues are labeled according to the Chothia numbering scheme. **(D)** Close-up view of key epitope residues (W241 and S274) at the CD38-daratumumab interface. The residues are shown as sticks, with neighboring CD38 residues (green) and daratumumab residues (light blue and orange) are shown as lines. Antibody residues are labeled according to the Chothia numbering scheme. **(E)** Flow cytometry histograms of CD38 WT or variants overexpressed in DF-1 cells. Staining using daratumumab or isatuximab combined with HB-7. For (E) n = 1-2 independent experiments for each group were performed.

As with CD20, we validated daratumumab and isatuximab binding to CD38^WT^ and CD38 carrying the different SNVs. As control antibody we used clone HB-7. CD38^WT^ expressing cells were double stained for both, daratumumab and control as well as isatuximab and control, respectively (Fig. 4E). In contrast, CD38^W241S^ and CD38^S274F^ affected daratumumab staining but neither control antibody nor isatuximab staining (Fig. 4E). CD38^S274F^ completely abolished daratumumab staining while CD38^W241S^ reduced staining almost completely. Thus, CD38^S274F^ and CD38^W241S^ will likely be completely resistant to daratumumab. On the other hand, CD38^G113R^ partially affected isatuximab staining but did not affect daratumumab binding. Therefore, isatuximab could be partially resistant to CD38^G113R^, but should be unaffected by all of the other identified SNVs.

### Impact of natural PD-1 variants on «checkpoint inhibitors»

The therapeutic antibodies targeting the B cell lineage targets CD20 and CD38 are cell depleters. Next, we analyzed frequently used “checkpoint inhibitors”. These antibodies do not directly target tumor cells but rather block an inhibitory signal provided by tumor cells that engage e.g. CTLA-4 or PD-1 (CD279) on T cells. As a result, cytotoxic T cells become dysfunctional and lose their ability to kill the cancer cells. Blocking the PD-1 mediated signal can restore T cell function resulting in tumor elimination (*12, 42–44*). We identified multiple SNVs at different PD-1 residues that might affect nivolumab or pembrolizumab binding. Among these, variants at residues S87, P89 and G90 are predicted to have the most significant impact on pembrolizumab binding, while variants at residues P28 are expected to affect nivolumab binding, although the effects are predicted to be less pronounced (Fig. 5A). Notably, S87 and P89 are located on a loop that serves as a primary anchor point for pembrolizumab, positioned within a groove at the interface of the heavy and light chain CDR loops (Fig. 5C). Conversely, P28 primarily interacts with W52 of the nivolumab heavy chain (Fig. 5, D and E). The following variants, all predicted to be neutral on the antigen, were selected for testing: CD279^P28L^, CD279^P28S^, CD279^S87R^, CD279^P89L^, CD279^P89R^ and CD279^P89T^.

**Fig. 5.**
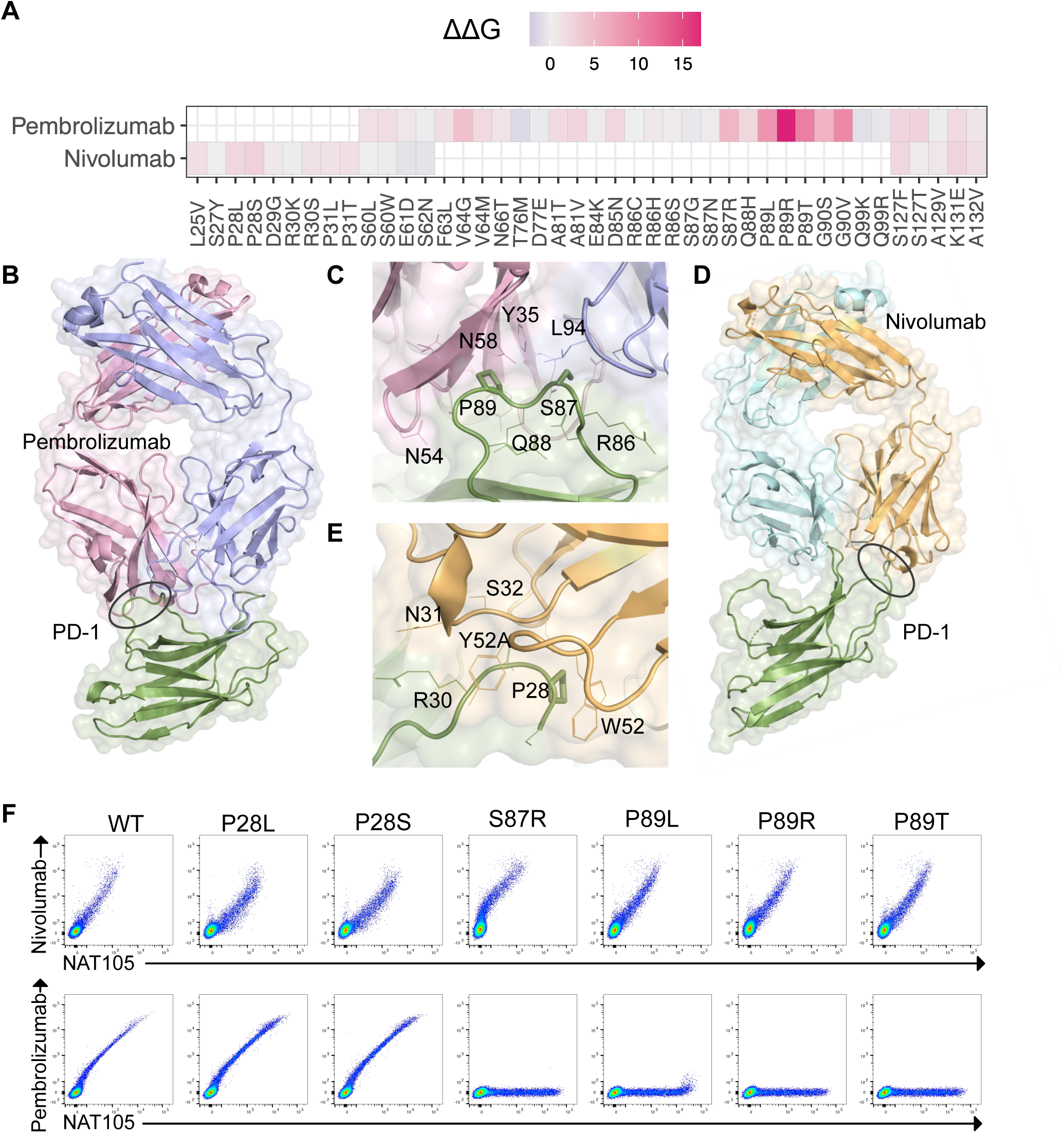
Impact of PD-1 epitope variants on the binding of pembrolizumab and nivolumab antibodies. **(A)** Predicted changes in binding free energy (ΔΔG) for missense epitope variants in the PD-1-pembrolizumab and PD-1-nivolumab complexes, represented as a heatmap. **(B)** Three-dimensional structure of the PD-1-pembrolizumab complex (PDB ID: 5GGS). The PD-1 antigen is shown in green, with the pembrolizumab heavy chain in pink and light chain in blue. Ovals indicate the locations of SNV sites at the interface of the complex. **(C)** Close-up view of key epitope residues (P89 and S87) at the PD-1-pembrolizumab interface. The residues are shown as sticks with neighboring PD-1 residues (green) and Pembrolizumab residues (pink and blue) shown as lines. Antibody residues are labeled according to the Chothia numbering scheme. **(D)** Three-dimensional structure of the PD-1-nivolumab complex (PDB ID: 5GGR). The PD-1 antigen is shown in green, with the nivolumab heavy chain in light blue and light chain in orange. Ovals indicate the locations of SNV sites at the interface of the complex. **(E)** Close-up view of the P28 residue at the PD-1-nivolumab interface. Residue P28 is depicted as a stick, with neighboring PD-1 residues (green) and nivolumab residues (light blue and orange) shown as lines. Antibody residues are labeled according to the Chothia numbering scheme. **(F)** Flow cytometry histograms of PD-1 WT or variants overexpressed in DF-1 cells. Staining using pembrolizumab or nivolumab combined with NAT105. For (F) n = 2-3 independent experiments for each group were performed.

Consistent with the computational prediction, all mutants stained for the control antibody NAT105 (Fig. 5F). Neither CD279^P28L^ nor CD279^P28S^ affected binding of nivolumab or pembrolizumab. In contrast, CD279^S87R^, CD279^P89L^, CD279^P89R^ and CD279^P89T^ strongly affected pembrolizumab but not nivolumab (Fig. 5F). CD279^S87R^, CD279^P89R^ and CD279^P89T^ completely abolished pembrolizumab binding while CD279^P89L^ displayed minimal residual binding. Thus, we found four SNVs at two different residues which nearly completely prevent pembrolizumab binding but none affecting nivolumab. Thus, as for the B cell targets CD20 and CD38, these CD279 SNVs prevent checkpoint inhibitor binding.

### Genetic variation affecting antibodies that bind epithelial cells

Finally, we analyzed ErbB2, a target found in epithelial tissues. About 20% of breast cancers and some ovarian and gastric cancers, i.e. solid tumors, overexpress ErbB2, providing the tumor cells a growth advantage (*13*). Since ErbB2 overexpression is associated with a poorer prognosis, trastuzumab was generated as a first generation ErbB2 targeting mAb. Trastuzumab is reported to exert its therapeutic effects through a dual mechanism, blocking ligand-independent dimerization and signaling (*45*), and mediating ADCC (*13, 46, 47*). Although trastuzumab combined with chemotherapy resulted in a significant clinical benefit and represented a major medical advance (*11*), second generation trastuzumab-based drugs were developed as ADCs: trastuzumab emtansine and trastuzumab deruxtecan. Both are based on the identical mAb (trastuzumab) but differ in the linker, toxic payload and drug-antibody ratio (*10, 48*). Trastuzumab deruxtecan is more potent in breast cancer (*48*) and approved for breast cancer and other ErbB2 expressing cancers including gastric, some forms of lung and some other cancers. Thus, trastuzumab-based drugs are very important in oncology.

Trastuzumab binds the juxtamembrane region of ErbB2 (domain IV) while pertuzumab binds to domain II (*45*), sterically blocking ligand-dependent dimerization and signaling (*49*) (Fig. 6, A and B), resulting in synergistic effects when administered together (*50*). We identified several SNVs across both epitopes (Fig. 6A), with ErbB2^P594H^ having the most significant effect on trastuzumab binding (Fig 6, A). Other identified variants are predicted to have minimal or no impact on antibody binding, except for ErbB2^G309E^, which is predicted to be pathogenic by AM and to impact pertuzumab binding (Fig. 6, A and C). G309 is located on a small turn that connects two β-strands and primarily participates in van der Waals interactions (Fig. 6C). Conversely, P594 is surrounded by aromatic side chains, including trastuzumab H3 residues Y100A and W95, mediating both intramolecular and intermolecular stacking interactions (Fig. 6D). To validate the computational analysis for ErbB2, we first tested trastuzumab and pertuzumab (biosimilars) binding without a linker-toxin payload and used clone 24D2 as a control antibody. ErbB2^WT^ cells bound all antibodies but ErbB2^G309E^ mildly affected pertuzumab but not trastuzumab binding (Fig. 6, E and F). Vice versa, ErbB2^P594H^ completely abolished trastuzumab binding but did not affect pertuzumab binding (Fig. 6, E and F). Furthermore, ErbB2^P594H^ appeared to slightly reduce ErbB2 expression. Thus, we experimentally validated that the newly described ErbB2^P594H^ SNV prevents trastuzumab binding.

**Fig. 6.**
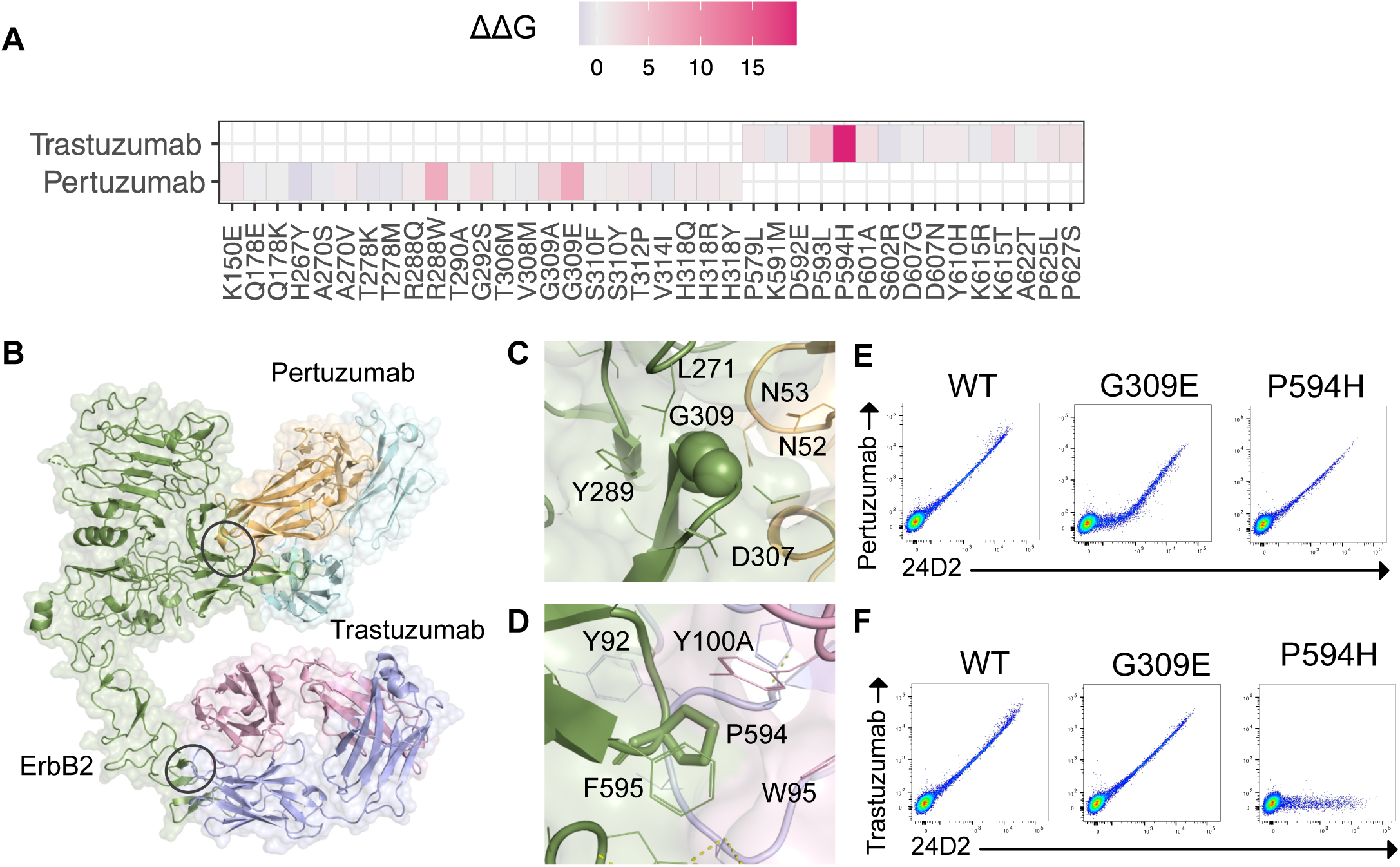
Impact of ErbB2 epitope variants on the binding of trastuzumab and pertuzumab antibodies. **(A)** Predicted changes in binding free energy (ΔΔG) for missense epitope variants in the ErbB2-trastuzumab and ErbB2-pertuzumab complexes, represented as a heatmap. **(B)** Three-dimensional structure representation of the ErbB2 ECD in complex with pertuzumab (PDB ID: 1S78) and trastuzumab (PDB ID: 1N8Z). The visualisation is derived from the structural superposition of ErbB2 (green). The pertuzumab heavy chain is shown light blue, and its light chain in orange. Trastuzumab heavy chain is shown in pink, and its light chain in blue. Ovals indicate the locations of SNV sites at the interface of the complex. **(C)** Close-up view of residue G309 at the ErbB2-pertuzumab interface. Residue G309 main chain atoms are depicted as spheres, with neighboring ErbB2 residues (green) and pertuzumab residues (light chain in orange) shown as lines. Antibody residues are labeled according to the Chothia numbering scheme. **(D)** Close-up view of residues P594 at the ErbB2-trastuzumab interface. Residue P594 is depicted as a stick, with neighboring ErbB2 residues (green) and trastuzumab residues (pink and blue) shown as lines. Antibody residues are labeled according to the Chothia numbering scheme. **(E)** Flow cytometry histograms of ErbB2 WT or variants overexpressed in DF-1 cells. Staining using pertuzumab combined with 24D2. **(F)** Flow cytometry histograms of ErbB2 WT or variants overexpressed in DF-1 cells. Staining using trastuzumab combined with 24D2. For (E) and (F) n = 2 independent experiments for each group were performed.

### A naturally occurring SNV confers resistance to clinical ADCs

Since we validated SNVs that negatively affected mAb binding for all tested antigens, we aimed to investigate SNV-mediated resistance to two highly potent and clinically very frequently used ADCs, trastuzumab emtansine and trastuzumab deruxtecan (*51*). For practical reasons, we mostly used research grade biosimilar antibodies in the FACS binding assays for experimental validation of the computational predictions. Rituximab was the exception for which we used the clinical biosimilar product (Rixathon). However, ADCs are very particular, even when the same mAb is used as a basis. Different linkers, toxins and using specific conjugation methods result in very different molecules with very specific properties (*10, 51*). Therefore, we wanted to test trastuzumab emtansine and trastuzumab deruxtecan with the original products Kadcyla and Enhertu, respectively. First, we analyzed ErbB2 expression on two breast cancer cell lines. ErbB2 expression is very low on MDA-MB-231 cells whereas the HCC1954 cell line overexpresses ErbB2 (Fig. 7A). We obtained both clinical ADCs from the hospital pharmacy and titrated trastuzumab emtansine (Kadcyla) and trastuzumab deruxtecan (Enhertu) on each cell line. Both ADCs did not kill the ErbB2 low cell line MDA-MB-231 at concentrations up to 2000pM. In contrast, HCC1954 cells were killed in a dose-dependent manner by both ADCs (Fig. 7B). To test resistance, we used CRISPR/Cas9-mediated homology directed repair (HDR) to engineer ErbB2^P594H^ into HCC1954 cells. Compared to control cells, HCC1954 electroporated with sgRNA/Cas9 ribonucleoprotein (RNP) resulted in ErbB2 KO as illustrated by a strong loss of trastuzumab and 24D2 binding (Fig. 7C). Addition of the HDR template encoding ErbB2^P594H^ resulted in a population of cells that stained for 24D2 but not trastuzumab which is indicative of the desired editing. To validate correct engineering, we purified (FACS sorted) “WT” (unedited), “KO” (Non homologous end joining = NHEJ) and knock-in (KI) (engineered) cells by flow cytometry. Next generation sequencing confirmed that the WT cells remained unedited, KO cells displayed indels whereas the vast majority of the purified KI cells contained the intended PAM mutation to increase editing efficiency and the CCC to CAC codon change resulting in ErbB2^P594H^ (Fig. 7D). We then tested susceptibility of WT, sorted WT, KO and KI HCC1954 cells to Kadcyla and Enhertu. While WT and sorted WT cells were killed by increasing concentrations of Kadcyla and Enhertu, the KO sorted (mainly ErbB2 KO) and KI sorted (mainly ErbB2^P594H^) cells were resistant to ADC-mediated killing (Fig. 7E). Thus, the newly identified ErbB2^P594H^ SNV, when engineered into a breast cancer cell line overexpressing ErbB2, resulted in complete resistance against the clinically used, highly potent drugs Kadcyla and Enhertu.

**Fig. 7.**
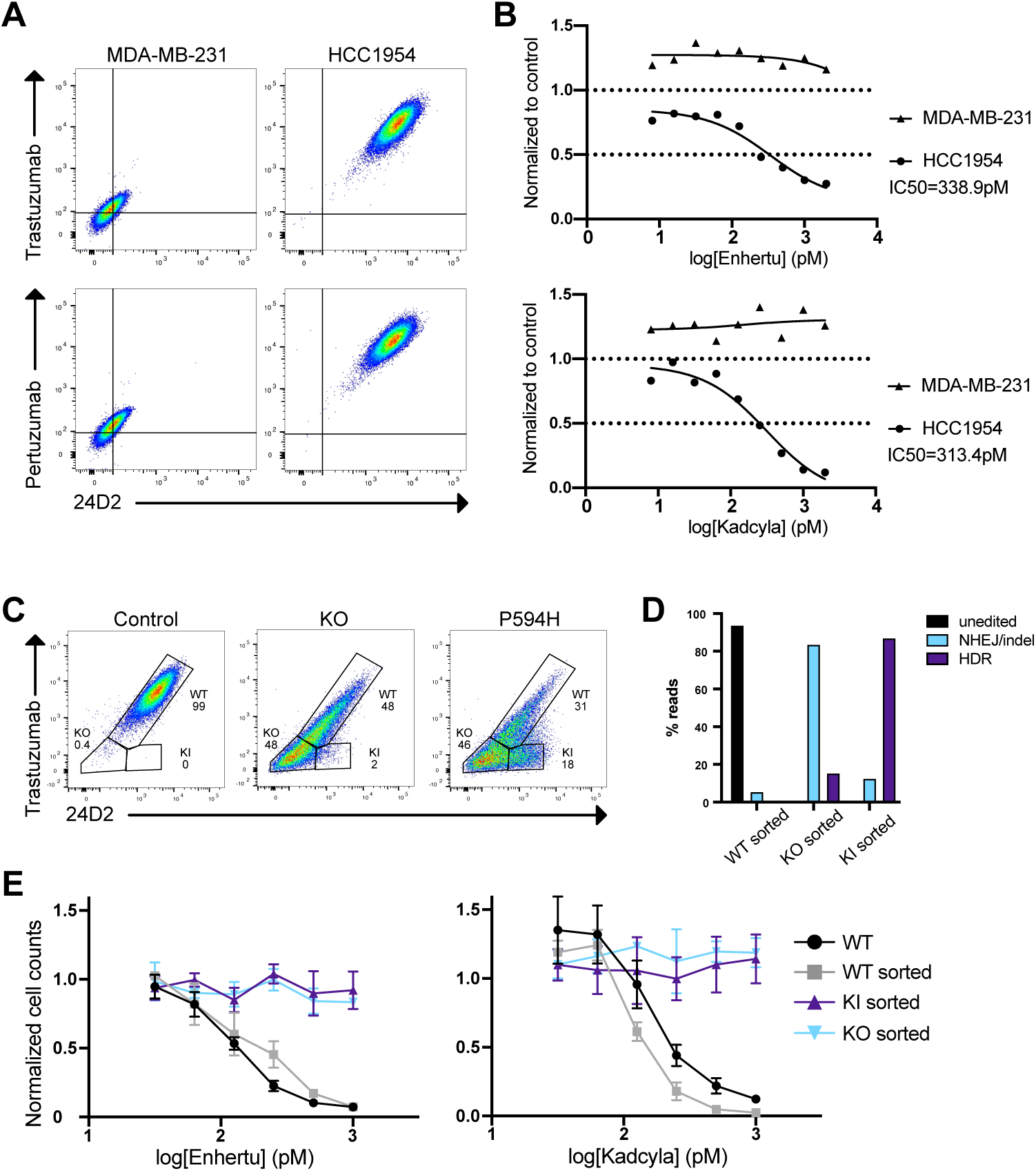
Engineering the natural SNV ErbB2^P594H^ in human breast tumor cells confers resistance to clinical ADCs. **(A)** Flow cytometry histograms showing ErbB2 expression in MDA-MB-231 and HCC1954 breast tumor cells lines. Staining was performed with ErbB2 antibodies recognising different epitopes (trastuzumab or pertuzumab combined with 24D2). **(B)** In vitro dose-titration (killing curves) of Enhertu or Kadcyla applied to MDA-MB-231 and HCC1954 cells. **(C)** Flow cytometry showing genome edited HCC1954 cells 3 days after electroporation. Staining with ErbB2 antibodies trastuzumab combined with 24D2. Control (cells electroporated with Cas9 protein only); KO (cells electroporated with Cas9 RNP); P594H (cells electroporated with Cas9 RNP and HDR template). In the flow cytometry histograms, double positive cells are defined as WT, double negative cells as KO and cells binding only to clone 24D2 as KI. **(D)** Amplicon-NGS sequencing of the targeted ErbB2 locus after sorting of control, KO and KI conditions. Unedited (WT codon); NHEJ/indel (indels at the Cas9 cutting site); HDR (KI codon and silent PAM mutation). **(E)** In vitro dose-titration (killing curves) of Enhertu or Kadcyla applied to edited and sorted HCC1954 and WT HCC1954 cells. For (A) n = 2 independent experiments for each group were performed. (B) and (E) n = 2-3 independent experiments in triplicates for each group were performed. In (E) data are mean +/-SD. (C) n = 3 independent experiments for each group were performed. For (D) n=1 experiment was performed.

## Discussion

Here, we analyzed how human genetic diversity in epitopes bound by therapeutic mAbs might affect antigen-specific therapies. We provide the first systematic study addressing the effect of human genetic variation on a given antibody. We focused on approved therapeutic antibodies and antibodies in clinical development. Notably, for every analyzed antibody we found multiple SNVs in their respective epitope regions. A substantial portion of these SNVs is predicted to alter antibody binding, either directly by disrupting the antibody-antigen interface without affecting the antigen’s structure or function, or indirectly, by altering the structure of the antigen itself. We experimentally validated the impact of 14 SNVs located at 11 epitope residues within 4 target proteins. For nine of these SNVs, we observed a significant reduction or complete loss of antibody binding, consistent with the computational predictions.

Our findings have significant implications for various stakeholders, including patients, physicians, healthcare systems, and the pharmaceutical industry. First and foremost, for patients carrying a specific SNV, treatment with a drug that cannot be effective means unnecessary procedures, exposure to unwarranted safety risks and false hope for therapeutic benefit. Although mAbs are generally well tolerated, the occurrence of infrequent but serious adverse events constitutes a concern. ADCs, on the other hand, are associated with treatment-related adverse events in ∼90% of patients, with serious events (grade ≥3) reported in 46% of cases on average (*10, 52*). Therefore, polymorphisms that could affect ADCs are particularly important. Frequently, ADC-related adverse events arise from target-independent effects associated with the intrinsic toxicity of the payload (*53, 54*), a key determinant of the maximum tolerated dose (*55*). Consequently, if the ADC fails to bind to its intended target due to a resistance-causing SNV, the entire dose is subject to potential target-independent, systemic toxicity, a highly undesirable outcome that should be avoided. Support for this comes from an ongoing clinical trial (NCT04849910) investigating patients with engineered ADC resistance (through the transplantation of CD33-deficient hematopoietic stem cells) and treated with a CD33-directed ADC (gemtuzumab ozogamicin (GO)). Compared to historic controls, low dose GO injections resulted in higher-than-expected ADC serum concentrations, comparable to healthy individuals carrying CD33-sufficient cells that received higher GO doses. By inference, applying the recommended GO dose - that was determined in a population of CD33-sufficient patients – would likely lead to higher GO serum concentrations and an increased risk for payload-related toxicity.

Physicians must be aware of the potential for resistance due to polymorphisms in the targeted epitopes. A diagnostic test for known epitope associated variants could help avoid ineffective treatments in patients carrying those variants and guide treatment decision, especially when multiple antibodies are available. For instance, CD20^N171Y^ results in resistance to rituximab but not ofatumumab (Fig. 3, F and G). Therefore, for CD20^N171Y^ carriers, ofatumumab should be chosen. Similarly, CD38^W241S^ and CD38^S274F^ are resistant to daratumumab but not isatuximab (Fig. 4E) (*30*). Therefore, a diagnostic test showing either of these variants would guide a physician to apply isatuximab rather than daratumumab. Vice versa, a patient carrying CD38^G113R^ should rather be treated with daratumumab than isatuximab. The same applies to other targets; for example, we identified 4 SNVs (S87R, P89L, P89R, P89T) affecting pembrolizumab but not nivolumab (Fig. 5F). Lastly, trastuzumab and pertuzumab also bind different epitopes on the ErbB2 antigen and are differentially affected by the ErbB2^P594H^ variant (Fig. 6, E and F). Therefore, ErbB2^P594H^ carriers should not be treated with trastuzumab-derived products but rather pertuzumab-based products.

Although the clinical significance of the newly identified variants in our study remains to be elucidated, some prior clinical evidence supports personalization of mAb-based therapies. A key consideration is the need to contextualize these findings within global and population-specific datasets, as some variants, though globally rare, may be more prevalent in certain populations. As an example, the C5^R885H^ polymorphism is associated with poor response to eculizumab, a treatment for paroxysmal nocturnal hemoglobinuria (PNH). In the overall European population, this variant is rare (GnomAD allele frequency: 2.5e−5), but in the East Asian population it reaches an allele frequency of 9.6e−3, with an estimated prevalence of approximately 3.5% in the Japanese population, similar among healthy individuals and PNH patients (*24*). The clinical relevance of this variant is well-documented. Among 345 treated PNH patients, there were 11 non-responders, all carrying the heterozygous C5^R885H^ variant (*24*). Similarly, a patient with hematopoietic stem cell transplantation-associated thrombotic microangiopathy who did not respond to eculizumab carried a C5^R885H^ variant. After switching to an alternative C5 inhibitor (Coversin), the patient initially improved, but ultimately died due to limited Coversin supply (*56*). This emphasises the importance of early identification of resistance-associated variants. Additional resistance variants, such as C5^R885C^ and C5^R885S^, have been linked to reduced response or lack of efficacy with eculizumab but switching to Coversin successfully treated a C5^R885S^ carrier patient (*57*). Like C5^R885H^, these and other variants are present in our dataset and are predicted to affect eculizumab binding or antigen function (Data File S2), suggesting that computational predictions and/or experimental diagnostic tests should be considered for personalization of mAb treatments. In general, variants influencing antibody binding or antigen function can result in a range of resistance phenotypes, ranging from mild binding affinity changes to complete loss of binding. These effects may be mitigated by compensatory factors such as optimized dosing or alternative therapies (*58*). Therefore, germline epitope variants may account for some non-responders to mAb therapies. In addition, secondary resistance could arise from somatic mutations acquired by the target cell as observed under CD19-directed CAR T therapy (*59*). In all cases, effective clinical management depends on early detection and characterization of resistance to enable timely and informed treatment decisions.

Limitations of our study and future directions: Although our computational analysis shows strong predictive value, predictions are not always accurate and require complementary evidence. For practical reasons we primarily validated predictions using research grade biosimilars, but we also confirmed several predictions with clinically approved commercial products, including the clinical biosimilar rituximab (Rixathon) and potent ADCs (Kadcyla, Enhertu). Furthermore, due to lack of availability, we did not analyze cells or tissues from patients with primary treatment failure. However, we provide many candidate SNVs that should be validated in future studies, ideally in patient cohorts where samples are available before and after treatment. Our results show that multiple high-impact variants can be identified for each antibody. However, precise quantification of the impact remains challenging. As computational methods and artificial intelligence (AI) continue to advance, they are expected to increasingly complement experimental approaches.

However, we argue that truly individualized predictions of drug response, based on patients’ genomic profiles, will require broader genetic variation coverage than is currently available to train accurate models. As noted, existing genetic studies often suffer from sparse data and population biases (*60*). This is not unlike the information needed to accurately predict genome engineering outcomes across populations. As genomic medicine and gene editing in medicine progress rapidly (*61*), human genetic diversity becomes increasingly relevant for medical therapies. Genetic variation can for instance affect the efficacy of CRISPR-based therapies or can generate off-target sites which can potentially affect the safety (*62–64*). For example, a common variant in individuals of African ancestry creates an off-target site for the guideRNA used in the first approved CRISPR-based therapy for sickle cell disease and transfusion-dependent ß-thalassemia (*65, 66*). In early studies the risk for this off-target effect was overlooked due to limited genomic diversity (*67*). Given the current bias toward European ancestry in genomic data (*60*), broader representation is crucial (*68*). Our findings underscore the need to capture global genetic diversity, as broader and more diverse datasets may reveal additional polymorphisms affecting antigen-specific therapies or identify populations at higher risk of therapy failure. In such a scenario, antigen-specific immunotherapies may be customized to specific populations, or even individuals. It is estimated that each individual’s genome differs from the reference human genome at 4-5 million sites, with over 99.9% of these differences being SNVs or short indels (*69*). Therefore, the variants annotated so far, and those analyzed here, represent only a fraction of the genetic variation that could impact antibody binding or treatment outcomes. As large-scale genomic datasets continue to expand, our ability to capture epitope diversity is expected to improve accordingly. On the other hand, the examples provided above along with findings from pharmacogenomic studies, highlight the presence of actionable variants in the majority of genotyped patients (*70*). As pharmacogenomics shifts from reactive testing of single genes to proactive multi-gene testing, pre-emptive testing may improve treatment efficacy, reduce adverse events, and lower costs for healthcare systems (*71, 72*). In line with this, recent FDA guidelines promote more comprehensive reporting of pharmacogenomic factors and initiatives to review gene-drug interactions (https://www.fda.gov/medical-devices/precision-medicine/table-pharmacogenetic-associations). In this context, our study addresses a gap in the investigation of variants affecting monoclonal antibody treatments, an area less explored compared to small molecule drugs.

Our findings have important implications for both clinical practice and therapeutic development. In particular, we argue that integrating genetic testing with accurate predictive models could help identifying patients less likely to benefit from specific therapies. This has the potential to reduce unnecessary treatments, minimize exposure to side effects, and optimize healthcare resources, ultimately improving patient care and outcomes. Moreover, our results may be valuable to the biotech and pharmaceutical industries. Accounting for genetic variation could help avoid false-negative outcomes that might otherwise bias clinical efficacy, and could inform the development of mAbs targeting alternative, non-overlapping epitopes, thereby enhancing treatment efficacy and coverage across genetically diverse patient groups.

## Acknowledgements

HCC1954 and MDA-MB-231 cell lines were a kind gift of Dr. M. Bentires-Alj. Clinical ADCs were a kind gift of Dr. Ott and the Pharmacy of the University Hospital Basel. We acknowledge the flow cytometry facility at the University of Basel and the Department of Biomedicine for support. Calculations were performed at sciCORE (http://scicore.unibas.ch/) scientific computing center at University of Basel. This project has received funding from the European Research Council (ERC) under the European Union’s Horizon 2020 research and innovation programme (grant agreement 818806 to L.T.J.) and institutional funds by the Department of Biomedicine. TS and EA acknowledge support by the SIB Swiss Institute of Bioinformatics and the Department Biozentrum of the University of Basel.

## Materials & Methods

### Study Design

Antibody specificity is a critical determinant of therapeutic efficacy and safety. Emerging evidence indicates that even single amino acid substitutions can significantly alter antibody-antigen recognition. In this study, we investigated the impact of naturally occurring missense variants on antibody-antigen interactions and their potential role in driving resistance to antigen-specific therapies. Publicly available databases were accessed to retrieve antibody sequences, structural information on target antigens and therapeutic antibodies, as well as human genetic variants associated with the antigens. Computational analyses were then performed to estimate the potential effects of these variants on the stability of the antigens and antibody-antigen complexes. For experimental validation, representative examples were selected to include a diverse set of antigens expressed on both hematologic and non-hematologic cells. We chose antibody therapeutics with different mechanism of action and with established clinical and commercial relevance. Computational predictions and experimental validation were done by different teams. Members of the experimental team were blinded for the FACS expression experiments but not the killing experiment.

### Dataset of therapeutic antibodies

Therapeutic antibody sequences were retrieved from the Thera-SabDab database (*73*), which catalogs antibody- and nanobody-based therapeutics recognized by the World Health Organization (WHO). Antibodies targeting human proteins with at least one experimental structure of the antibody-antigen complex were selected. The corresponding PDB files were renumbered using the Chothia scheme (*74*) by aligning heavy and light chain sequences to isotype-specific Hidden Markov Model (HMM) profiles (*75*). The OpenStructure framework (*76*) was used to assess sequence identity between therapeutic sequences and the amino acid sequences of structural data, retaining only cases with 100% sequence identity. To exclude artifacts and distinguish biological interactions from crystallographic contacts, only structures with antibody-antigen interactions mediated by complementarity determining regions (CDRs) were retained. Antibodies with identical variable heavy and light chain amino acid sequences were clustered together and a single representative was selected for analysis. The full list of antibodies analysed in this study is provided in Data file S1. Clinical trial status and therapeutic indications were manually curated using DrugBank (*77*), ClinicalTrials.gov, and the literature.

### Human single nucleotide variants

The UniProt IDs of the protein targets (*78*) were used to query the European Bioinformatics Institute (EBI) database via API (*79*) to retrieve a comprehensive collection of human genetic missense variants aggregated from multiple sources, including the Exome Aggregation Consortium (ExAC) (*80*), gnomAD (*81*), the 1000 Genomes Project (*69*), TopMed (*82*), and ClinVar (*83*), among others, along with disease annotation for the identified variants. Data was retrieved on October 2022. From this dataset, only missense variants were retained for further analysis.

### Computational analysis of SNV-Associated amino acid variants

Variants were mapped onto the available 3D structures of antibody-antigen complexes using the Var3D variant annotation pipeline available at https://git.scicore.unibas.ch/schwede/var3d. Epitopes were defined as the set of antigen residues within 5Å to any residue of the antibody in the 3D complex. Sequence- and structure-related features for SNV-associated amino acid sites were computed by Var3D using the OpenStructure implementation (*76*). Specifically, per-residue relative solvent accessibility (RSA) was computed using the NACCESS implementation of the Lee & Richards method (*84*). For each target sequence, a multiple sequence alignment (MSA) was generated based on a Jackhammer search (e-value < 1e−4) against the UniRef90 database (Release 2021_04) (*85, 86*). The resulting MSAs were used to compute Shannon entropy (doi: 10.1002/j.1538-7305.1948.tb01338.x) and ConSurf conservation scores. ΔΔG values for each amino acid variant were computed using FoldX (*27*), with the 3D structure of the antibody-antigen complex and the isolated antigen structure as inputs. AlphaMissense (*28*) predictions were retrieved from https://console.cloud.google.com/storage/browser/dm_alphamissense.

### Eukaryotic cell lines

UMNSAH/DF-1 cells were purchased from ATCC (Cat. CRL-12203) and were expanded in Dulbecco’s Modified Eagle’s Medium; high glucose (4.5g/l) (Sigma-Aldrich), supplemented with 10% FCS (Gibco Life Technologies) and 2 mM GlutaMAX (Gibco) at a temperature of 39°C. HCC1954 cells were a gift from M. Bentires-Alj (Department of Biomedicine, Basel, Switzerland) and were expanded in RPMI-1640 media (Sigma-Aldrich) supplemented with 10% heat-inactivated FCS (Gibco Life Technologies), 2 mM GlutaMAX. MBA-MB231 cells were a gift from M. Bentires-Alj and were expanded in Dulbecco’s Modified Eagle’s Medium; high glucose (4.5g/l) (Sigma-Aldrich), supplemented with 10% heat-inactivated FCS, and 2 mM GlutaMAX.

### Cloning and expression of various recombinant WT human genes and their variants

Full-length cDNAs of human CD20 (NM_021950.3), human CD38 (NM_001775.2), human CD279 (NM_005018.2) and human CD340 (NM_004448.2) were obtained in a pCMV3 vector (Cat# HG11007-UT, Cat# HG10818-UT, Cat.HG10377-UT, Cat# HG10004-UT; Sino Biological). The hygromycin sequence was replaced by a neomycin resistance cassette by restriction digestion. The SNVs of the human CD20, CD38, CD279 and CD340 variants selected for experimental validation were introduced into the cDNAs using the Q5^®^ Site-Directed Mutagenesis Kit (New England Biolabs, Cat. E055AS) (see table “Table S1”). 1 × 10^6^ UMNSAH/DF-1 cells were plated the day before transfection in a well of a 6 well plate. The following day the cells were transfected with 6.5µg DNA of the pCMV3 vector encoding WT cDNA or its variants and 19.5µg polyethyleneimine (PEI, Cat#23966; Polysciences) in 200µl of serum free medium and incubated for 2 days at 39°C before further analysis.

### Engineering of HCC1954 cell line

1 × 10^6^ HCC1954 cells were resuspended in 20 μl Lonza-supplemented SF EP buffer. Cas9 RNPs were freshly prepared as described prior to each EP (*20*). Thawed crRNA and tracrRNA (purchased from IDT Technologies; at 200 μM) were mixed in a 1:1 M ratio (120 pmol each), denatured at 95°C for 5 min, and annealed at room temperature (RT) for 10 min to complex an 80 μM gRNA solution. Polyglutamic acid (15–50 kD at 100 mg/ml; Sigma-Aldrich) was added to the gRNA in a 0.8 : 1 volume ratio. To complex RNPs, 60 pmol recombinant Cas9 (University of California Berkeley; at 40 μM) was mixed with the gRNA (molar ratio Cas9:gRNA = 1:2) and incubated for 20 min at RT. HDRT (purchased from IDT Technologies; 50 pmol) and RNPs (60 pmol) were mixed and incubated for 5 min. The cells were added to the mix and the total volume was transferred to 16-well Nucleocuvette Strips. Electroporation was performed with the 4D-Nucleofector system (Lonza) with Program FF-150. Immediately following EP, 80 μl of prewarmed supplemented medium was added to each cuvette and incubated at 37°C. After 20 min, the cells were transferred into 6-well culture plates containing 2 ml of media. The HDRT contained the desired SNV and an additional silent mutation disrupting the PAM sequence to increase HDR efficiency. For crRNA and HDRT sequences see Table S2. Cells were cultured and expanded for FACS analysis and cell sorting.

### Flow cytometry and cell sorting

Unlabeled antibodies obtained from ProteoGenix and the Hospital Pharmacy, were labeled using Alexa Fluor™ 647 NHS Ester or Alexa Fluor™ 488 NHS Ester, both from ThermoFisher Scientific. Antibodies used for flow cytometry can be found in the Table S3. Flow cytometry was performed on BD LSRFortessa with the BD FACSDiva Software and the data were analyzed with FlowJo Software. For cell sorting, the HCC1954 engineered cells were pelleted, stained with ErbB2 antibodies (clone 24D2 and trastuzumab) and resuspended in FACS buffer (PBS + 2% FCS). Sorting was performed on BD FACSAria.

### Genomic DNA extraction and next generation sequencing (NGS)

Genomic DNA extraction was performed using QuickExtract (Lucigen) and the genomic DNA concentration was measured with a Qubit device (ThermoFisher). For NGS, a targeted amplicon library was prepared as three-step PCR protocol as described and paired-end sequenced on an Illumina Miniseq instrument using the Illumina Miniseq Mid output kit (300 cycles) with 50% PhiX spike-in (illumina) (*19*). The ErbB2 locus was amplified using the primers listed in Table S1. After demultiplexing, each sample was assessed for quality and processed using CRISPResso2 (*87*). Quantification of the different alleles (WT, KO and KI) was performed using a custom script as previously described (*19*).

### Killing assay

20’000 HCC1954 cells (WT, KO and KI) were plated in 96-well plates in medium with corresponding concentrations of Kadcyla or Enhertu. Following a 72 h incubation period, cells were collected and stained for viability and with ErbB2 antibodies (clone 24D2 and trastuzumab). Data were acquired on BD LSRFortessa.

## Author contributions

Conceptualization: LTJ, RL

Methodology: RM, EA, GC, TS, LTJ, RL

Software: EA, RL

Investigation: RM, EA, GC, RL

Visualization: RM, EA, RL

Funding acquisition: EA, TS, LTJ

Project administration: RM, LTJ, RL

Supervision: RM, TS, LTJ, RL

Writing – original draft: LTJ, RL

Writing – review & editing: RM, EA, GC, TS, LTJ, RL

## Data and materials availability

All data are available in the main text or the supplementary materials.

## Competing interests

RM, TS, LTJ, RL: Co-founder, holding equity of Cimeio Therapeutics AG (Cimeio). Inventor on granted patents and patent applications related to immune cell engineering. LTJ: Cimeio board member. Sponsored research agreement with Cimeio. Received speaker fees from Novartis. Paid consultant for Kyowa Kirin.

All other authors declare that they have no competing interests.

---

**Supplementary Figure S1.**
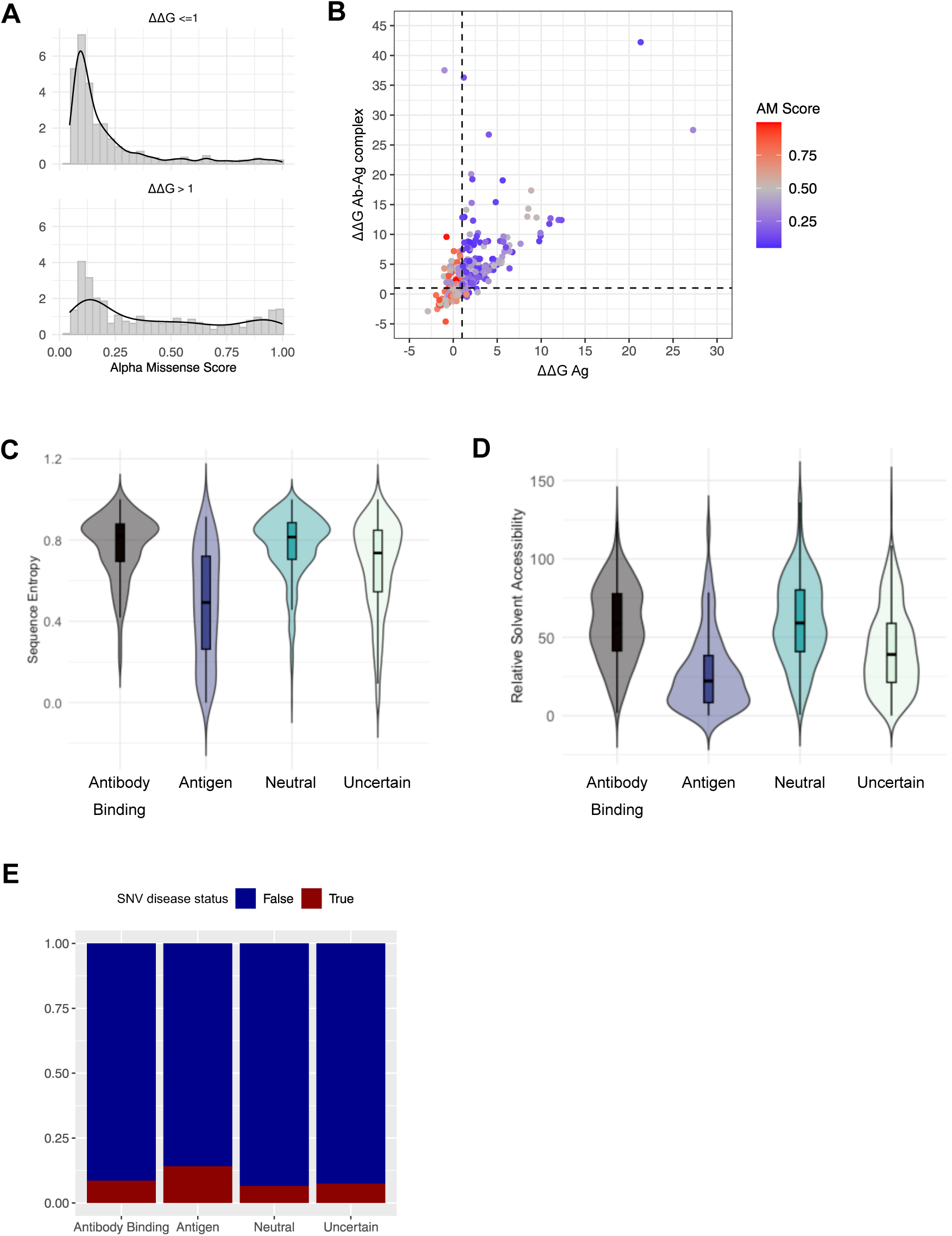
(**A**) Distribution of AlphaMissense (AM) scores for variants predicted with ΔΔG ≤ 1 and ΔΔG > 1. (**B**) Predicted effects (ΔΔG) of missense epitope variants on antibody-antigen complexes (y-axis) versus antigens alone (x-axis). Data points are color-coded based on AM scores. Variants with low AM scores (likely benign) are shown in blue, high scores (likely pathogenic) are shown in red, and intermediate scores (ambiguous) are shown in grey. (**C**) Distribution of sequence entropy across antigen sites associated with the four SNV classes. (**D**) Distribution of relative solvent accessibility across antigen sites associated with the four SNV classes. (**E**) Distribution of disease-associated variants across the four SNV classes.

**Table S1.**
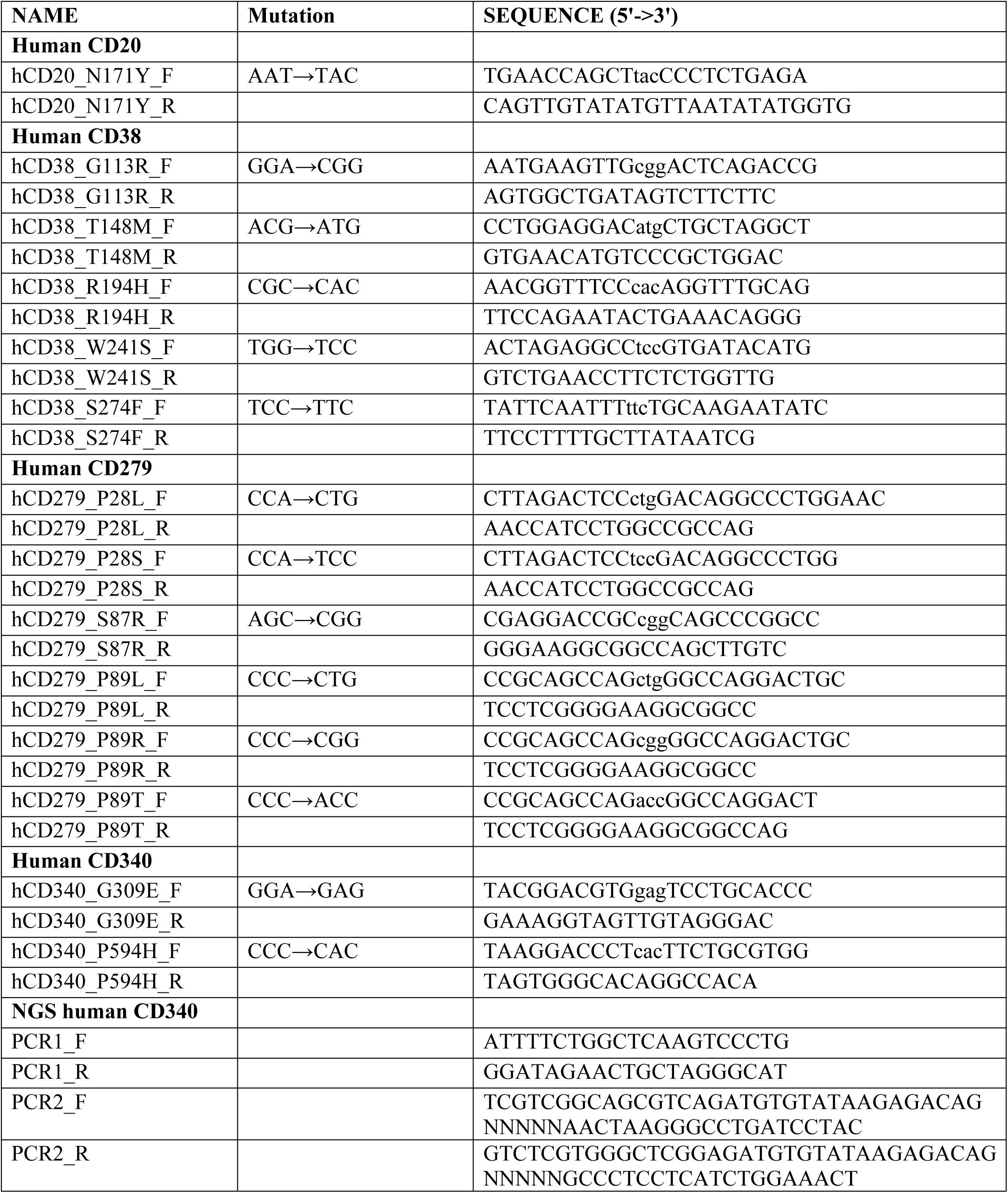
Sequences of the primers used to introduce the mutations using the Q5^®^ Site-Directed mutagenesis kit and NGS primers. F = Forward; R = Reverse.

**Table S2.**
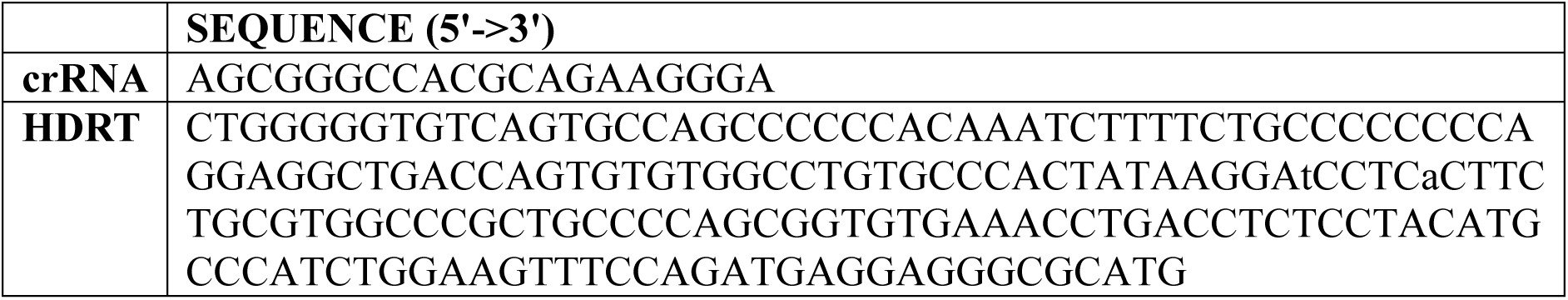
Sequences of crRNA and HDRT used to introduce the ErbB2^P594H^ variant into the HCC1954 cell line.

**Table S3.** List of antibodies used for the FACS stainings. Unlabeled antibodies obtained from ProteoGenix and the Hospital Pharmacy were labeled in house using Alexa Fluor™ 647 NHS Ester (Catalogue No. A20006) or Alexa Fluor™ 488 NHS Ester (Catalogue No. A20000) both from ThermoFisher Scientific.

| **Target** | **Clone** | **Vendor** | **Catalogue No.** | **RRID** |
| --- | --- | --- | --- | --- |
| **CD20** | 2H7 | Biolegend | 980214 | AB_2941644 |
|  | Rixathon | Hospital Pharmacy (Sandoz) |  |  |
|  | Ofatumumab | ProteoGenix | PX-TA1146-1MG |  |
| **CD38** | HB-7 | Biolegend | 356603 | AB_2561899 |
|  | HIT-2 | Biolegend | 980302 | AB_2616620 |
|  | S17015A | Biolegend | 397103 | AB_2810610 |
|  | S17015F | Biolegend | 397203 | AB_2810612 |
|  | Daratumumab Biosimilar | ProteoGenix | PX-TA1213-1MG | - |
|  | Isatuximab Biosimilar | ProteoGenix | PX-TA1377-1MG | - |
| **CD279** | NAT105 | Biolegend | 367403 | AB_2566064 |
|  | EH12.2H7 | Biolegend | 329905 | AB_940481 |
|  | Nivolumab Biosimilar | ProteoGenix | PX-TA1004-1MG | - |
|  | Pembrolizumab Biosimilar | ProteoGenix | PX-TA1007-1MG | - |
| **CD340** | 24D2 | Biolegend | 324406 | AB_756122 |
|  | Trastuzumab | ProteoGenix | PX-TA1005-1MG | - |
|  | Pertuzumab Biosimilar | ProteoGenix | PX-TA1058-1MG | - |
|  | Enhertu | Hospital Pharmacy (Daiichi Sankyo) |  |  |
|  | Kadcyla | Hospital Pharmacy (Roche) |  |  |
| **FcR Blocking Reagent, human** |  | Miltenyi | 130-059-901 | AB_2892112 |
| **Zombie UV Fixable Viability Kit** |  | BioLegend | 423108 | - |

